# The myosin regulatory light chain Myl5 localizes to mitotic spindle poles and is required for proper cell division

**DOI:** 10.1101/2020.06.04.134734

**Authors:** Ivan Ramirez, Ankur A. Gholkar, Erick F. Velasquez, Xiao Guo, Jorge Z. Torres

## Abstract

Myosins are ATP-dependent actin-based molecular motors critical for diverse cellular processes like intracellular trafficking, cell motility and cell invasion. During cell division, myosin MYO10 is important for proper mitotic spindle assembly, the anchoring of the spindle to the cortex, and positioning of the spindle to the cell mid-plane, while myosin MYO2 functions in actomyosin ring contraction to promote cytokinesis. However, myosins are regulated by myosin regulatory light chains (RLCs), and whether RLCs are important for cell division has remained unexplored. Here, we have determined that the previously uncharacterized myosin RLC Myl5 associates with the mitotic spindle and is required for cell division. Myl5 localized to the mitotic spindle poles and spindle microtubules during early mitosis, an area overlapping with MYO10 localization. Depletion of Myl5 led to defects in chromosome congression and to a slower progression through mitosis. We propose that Myl5 is a novel myosin RLC that is important for cell division.

## INTRODUCTION

The proper assembly of the bipolar mitotic microtubule spindle is critical to the fidelity of chromosome congression and segregation during cell division^1^. During development, the anchoring and positioning of the mitotic spindle regulates the establishment of the cell division plane that is critical for cell fate determination^2^. Important to mitotic spindle anchoring and positioning are astral microtubules that radiate out from the spindle poles and make contacts with the cell cortex^2–4^. Abnormal spindle assembly and orientation can result in defective cell divisions that can lead to developmental and proliferative diseases^2–4^. Although numerous components involved in assembling and orienting the spindle are known^1, 2^, the full complement of factors and the molecular signaling pathways that govern these events are not completely understood. Increasing evidence indicates that both the actin and microtubule cytoskeletal systems are necessary for proper cell division^5^. Although actin has been highly studied within the context of interphase cells where it establishes the cellular architecture and regulates numerous important processes like cell motility, intracellular trafficking, cell signaling pathways and gene expression^6–9^, less is known about its role during early cell division. However, actin has been shown to be critical for anchoring the spindle through microtubule-actin interactions at the cell cortex, for spindle positioning at the mid plane, and for actomyosin cellular constriction during cytokinesis^5, 10–12^. Additionally, evidence indicates that an actin mesh assembly supports the bipolar meiotic and mitotic spindles, where actin provides rigidity and aids in focusing the spindle^13–15^. Therefore, actin plays an important role in ensuring the fidelity of cell division.

Myosins are ATP-dependent actin-based molecular motors that perform a variety of functions in muscle contraction, cargo transport, cell adhesion, and cell division; including spindle assembly, spindle orientation and cytokinesis^16, 17^. During cell division, the unconventional myosin-2 (MYO2) is critical for acto-myosin ring contraction during cytokinesis, which is essential for bisecting one cell into two daughter cells^18, 19^. Of interest, the unconventional myosin-10 (MYO10) has been shown to be an important factor for the architecture and function of the mitotic spindle through its binding to both actin and microtubules^13, 14, 20, 21^. MYO10 localizes to the spindle poles throughout mitosis and depletion of MYO10 leads to structural defects in the mitotic microtubule spindle, chromosome congression defects, and chromosome segregation defects^13, 14, 21^. Therefore, myosins perform important functions that are necessary for a productive cell division. The unconventional myosin holoenzymes typically consists of heavy and light chains^16^. Myosin light chains are required for the structural integrity of the myosin holoenzyme and have regulatory functions on the activity of the protein complex^16, 22, 23^. There are two major groups of myosin light chains, the Essential Light Chains (ELCs) and the Regulatory Light Chains (RLCs)^22, 23^. The ELCs are essential for the enzymatic activity of the myosin and removal or depletion of the ELCs from the myosin leads to a dramatic loss of myosin enzymatic activity^22, 23^. The RLCs are involved in regulating the enzymatic activity of the myosin and their removal or depletion typically leads to moderate effects on myosin activity^22, 23^. Although MYO2 and MYO10 have important roles in cell division, the role of RLCs (if any) in cell division has remained unexplored.

A recent genetic screen for novel cell division proteins identified myosin light chain 5 (Myl5) as a putative factor important to cell division (Torres lab unpublished). Although Myl5 has remained poorly characterized, based on its protein sequence similarity it is predicted to be a myosin RLC^24^. Dysregulation of Myl5 protein levels has been linked to tumorigenesis and metastasis in glioblastoma multiforme, cervical carcinoma and breast cancer^25–27^. For example, Myl5 gene expression is upregulated in late stage cervical cancer patients and is associated with poor survival^25^. Additionally, Myl5 overexpression promoted tumor cell metastasis in a cervical cancer mouse model^25^. Here, we have discovered that Myl5 is important for mitotic spindle assembly, chromosome congression, and proper cell division. Myl5 localizes to the spindle poles during mitosis, indicating that its localization is cell cycle phase dependent. Myl5 co-localized with spindle pole proteins and MYO10. Importantly, depletion of Myl5 led to spindle assembly defects and lagging chromosomes. While Myl5 overexpression led to a faster progression through cell division. These results indicate that Myl5 has important functions in spindle assembly and chromosome segregation and that it functions with MYO10 to perform these functions.

## RESULTS

### *In silico* analysis of Myl5

Our recent genetic RNAi screen for novel cell division proteins led us to discover myosin light chain 5 (Myl5), an uncharacterized hypothetical myosin regulatory light chain (RLC) of the MLC2 type. Human Myl5 is a 173 amino acid protein with 3 EF hand domains predicted to be important for calcium binding in other myosin regulatory light chains (**Fig. 1A**)^23, 28^. An OrthoDB^29^ ortholog analysis indicated that Myl5 was conserved among vertebrates (**Fig. 1B**). An Online Mendelian Inheritance in Man (OMIM) search showed that the *MYL5* gene was within the 4p16.3 region where the Huntington Disease locus is located. However, Myl5 has not been linked to inherited human diseases. Due to recent studies showing the dysregulation of Myl5 gene expression in breast^27^, cervical^25^, and brain cancers^26^, we sought to determine if Myl5 was widely dysregulated in other types of cancers. Interestingly, analysis of Myl5 using The Cancer Genome Atlas (TCGA) Pan-Cancer Atlas^30^ showed that Myl5 had an increased alteration (mutation; fusion; amplification; deletion) frequency in some types of tumors, especially stomach adenocarcinomas, ovarian epithelial tumors, and bladder urothelial carcinomas (**Fig. 1C**). Additionally, ovarian epithelial tumors, bladder urothelial carcinomas, sarcomas, adrenocortical carcinomas, and pancreatic adenocarcinomas displayed the highest frequencies of Myl5 amplification (**Fig. 1C**). Together, these analyses showed that the Myl5 protein is conserved among vertebrates and that the Myl5 gene is mutated and has altered expression patterns in numerous types of cancers.

**Figure 1.**
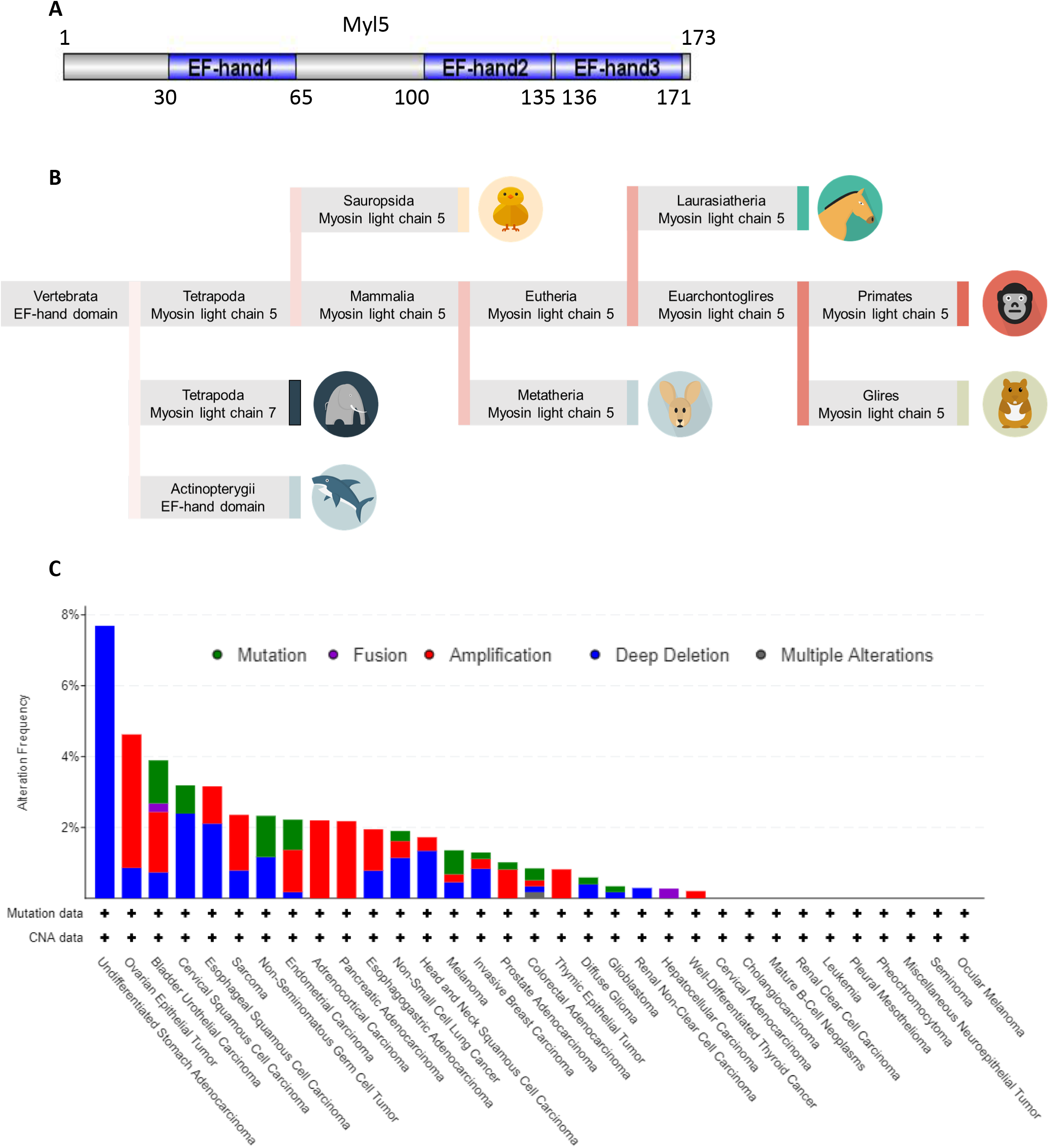
*In silico* analyses of Myl5. (**A**) Schematic of the human Myl5 (UniProtKB-Q02045) protein domain architecture with the EF hand domains highlighted in blue. The number of amino acid residues are indicated. (**B**) OrthoDB ortholog analysis showing that Myl5 is conserved among vertebrates. (**C**) Analysis of *MYL5* alteration frequencies in cancer using The Cancer Genome Atlas (TCGA) Pan-Cancer Atlas. Alteration frequency is on the y-axis and cancer type is on the x-axis. The type of alteration is color coded as indicated in the legend. Note that *MYL5* is frequently altered in cancer, especially amplifications in ovarian, bladder, adrenocortical and pancreatic cancers.

### Myl5 localizes to the spindle poles and spindle microtubules during cell division

Although previous genomic and bioinformatic studies had implicated Myl5 in myosin related functions and in tumorigenesis, its biological function had remained poorly characterized. To begin to understand the cellular role of Myl5 and its link to tumorigenesis, we analyzed its subcellular localization throughout the cell cycle. First, we generated a LAP(GFP-TEV-S-Peptide)-Myl5 inducible stable cell line that expressed GFP-Myl5 upon induction with Dox (**Fig. S1A**)^31, 32^. The LAP-Myl5 cell line was treated with Dox for 16 hours to express GFP-Myl5 and cells were fixed, stained with Hoechst 33342 DNA dye, and anti-a-Tubulin and anti-GFP antibodies and imaged by immunofluorescence microscopy. During interphase Myl5 was dispersed throughout the nucleus and cytoplasm of the cell (**Fig. 2A)**. Interestingly, Myl5 localized to the spindle poles in early mitosis, and to a lesser extent the mitotic spindle, and remained associated with the poles until mitotic exit (**Fig. 2A**). To further define the Myl5 subcellular localization in early mitosis, we performed immunofluorescence colocalization studies with centrosome and spindle pole markers. The Myl5 localization signal overlapped with NUMA at the spindle poles and encompassed the Pericentrin and Centrin signals, which stained the centrosomes (**Fig. 2B and C and Fig. S1B**). Furthermore, in cells with high levels of Myl5 expression, Myl5 also co-localized with TPX2 on the spindle microtubules (**Fig. S1C**). Consistent with the GFP-Myl5 mitotic localization, immunofluorescence microcopy of HeLa cells with anti-Myl5 antibodies showed that endogenous Myl5 also localized to the spindle poles and spindle microtubules during mitosis (**Fig. S2**). Due to the change in Myl5 localization at mitotic entry, we next asked if Myl5 protein levels were also cell cycle regulated. HeLa cells were synchronized in G1/S with thymidine treatment, released into the cell cycle, cells were harvested every hour, and protein extracts were prepared. Immunoblot analysis of these samples with anti-Myl5 and anti-Cyclin B antibodies indicated that endogenous Myl5 protein levels remained steady throughout the cell cycle and only a minor decrease was observed during mitotic exit, a time when mitotic Cyclin B levels decreased (**Fig. 2D**). Together, these results indicated that the Myl5 protein is abundant throughout the cell cycle and that it undergoes a dynamic cell cycle dependent change in subcellular localization where it redistributes from the cell cytoplasm in interphase to the spindle poles during mitotic entry and remains associated with the poles throughout mitosis.

**Figure 2.**
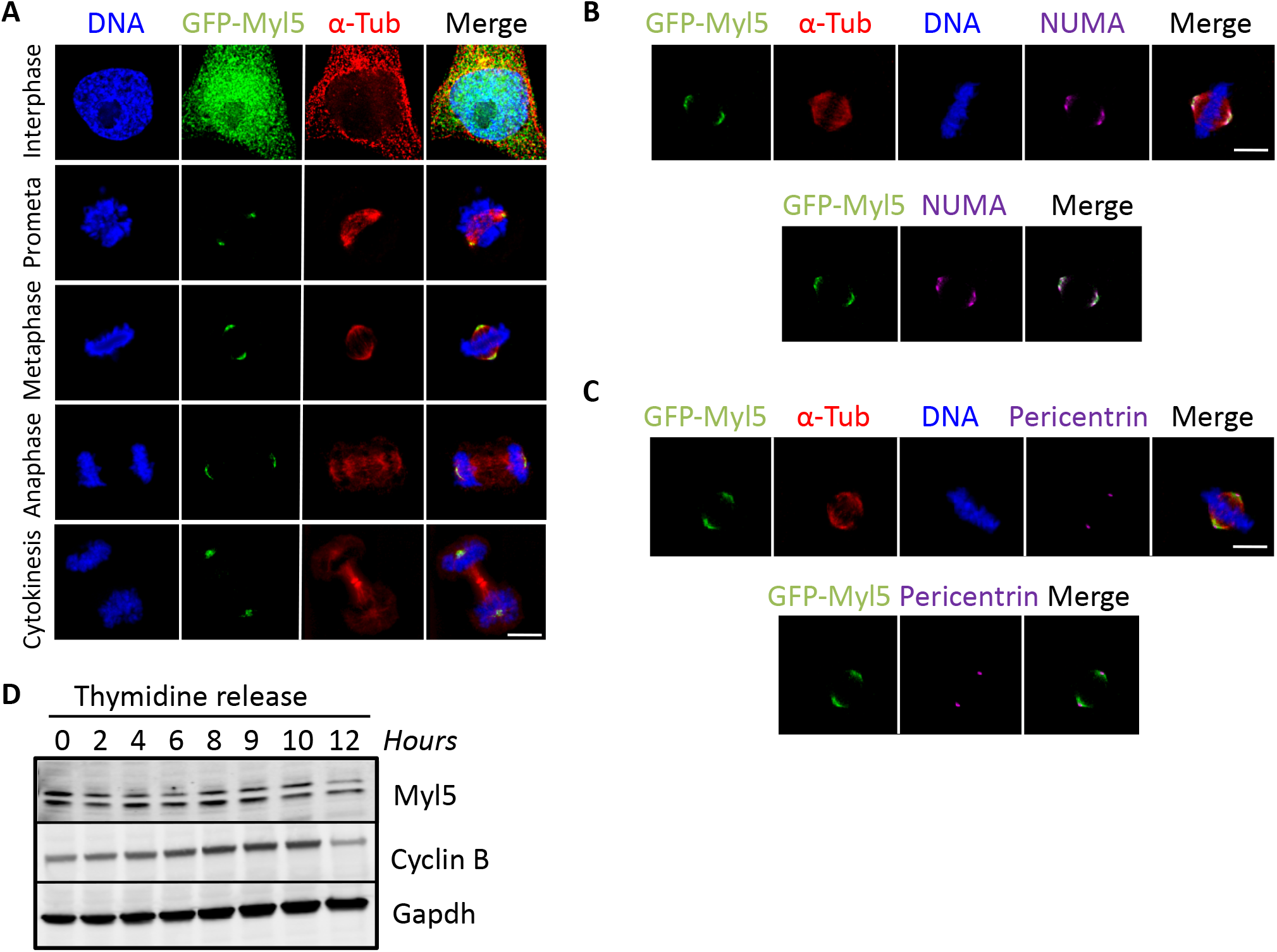
Myl5 localizes to mitotic spindle poles during mitosis. (**A**) The LAP (GFP-TEV-S-Peptide)-tagged-Myl5 HeLa inducible stable cell line was treated with Dox for 16 hours to express GFP-Myl5 and cells were fixed, stained with Hoechst 33342 DNA dye, and anti-a-Tubulin and anti-GFP antibodies and imaged by immunofluorescence microscopy. Images show the cell cycle subcellular localization of GFP-Myl5 during interphase, prometaphase, metaphase, anaphase and cytokinesis. Note that GFP-Myl5 localizes to the mitotic spindle poles during mitosis. Bar indicates 5μm. (**B-C**) Same as in **A**, except that cells were also stained with anti-NUMA (**B**) or anti-Pericentrin (**C**) antibodies. Note that GFP-Myl5 localizes to the mitotic spindle poles overlapping with the NUMA and Pericentrin localization signal. Bar indicates 5μm. (**D**) Analysis of endogenous Myl5 protein levels throughout the cell cycle. HeLa cells were synchronized in G1/S, released into the cell cycle and cells were harvested at the indicated time points. Protein extracts were prepared, resolved by SDS-PAGE, transferred to a PVDF membrane and immunoblotted with the indicated antibodies. Note that the Myl5 protein levels remain constant throughout the cell cycle, whereas the Cyclin B levels increase as the cells enter mitosis (6-10 hour time points). Gapdh is used a loading control.

### Myl5 is required for proper cell division

Next, we asked if Myl5 was required for cell division by depleting Myl5 in HeLa cells. First, we sought to identify siRNA oligonucleotides which reduced Myl5 protein levels to less than 10% compared to non-targeting control siRNA. HeLa cells were treated with non-targeting control siRNA (siCtrl) or siRNAs targeting Myl5 (siM1-siM4) for 72 hours and cell lysates were prepared and analyzed by immunoblotting. The siM1-siM4 oligonucleotides depleted Myl5 protein levels to undetectable levels (**Fig. 3A and Fig. S3A**). Next, we sought to analyze the consequences of depleting Myl5 protein levels during cell division. HeLa cells were treated with siCtrl or siM1-siM4 siRNAs for 72 hours. The cells were then fixed and co-stained with Hoechst 33342 (to visualize the DNA) and anti-a-Tubulin antibodies to detect the mitotic microtubule spindle. Interestingly, depletion of Myl5 led to a significant increase in cells with a defective mitosis, including an increase in the percentage of prometaphase cells with multipolar spindles (siM1= 20.75±3.59, p=.0004 compared to siCtrl= 7.5±1.29) and anaphase cells with lagging chromosomes (siM1= 32.25±2.5, p<.0001 compared to siCtrl= 8.25±3.1) (**Fig. 3B-E and Fig. S3B-F**). Together, these results indicated that Myl5 was required for proper cell division and that its depletion led to cell division errors.

**Figure 3.**
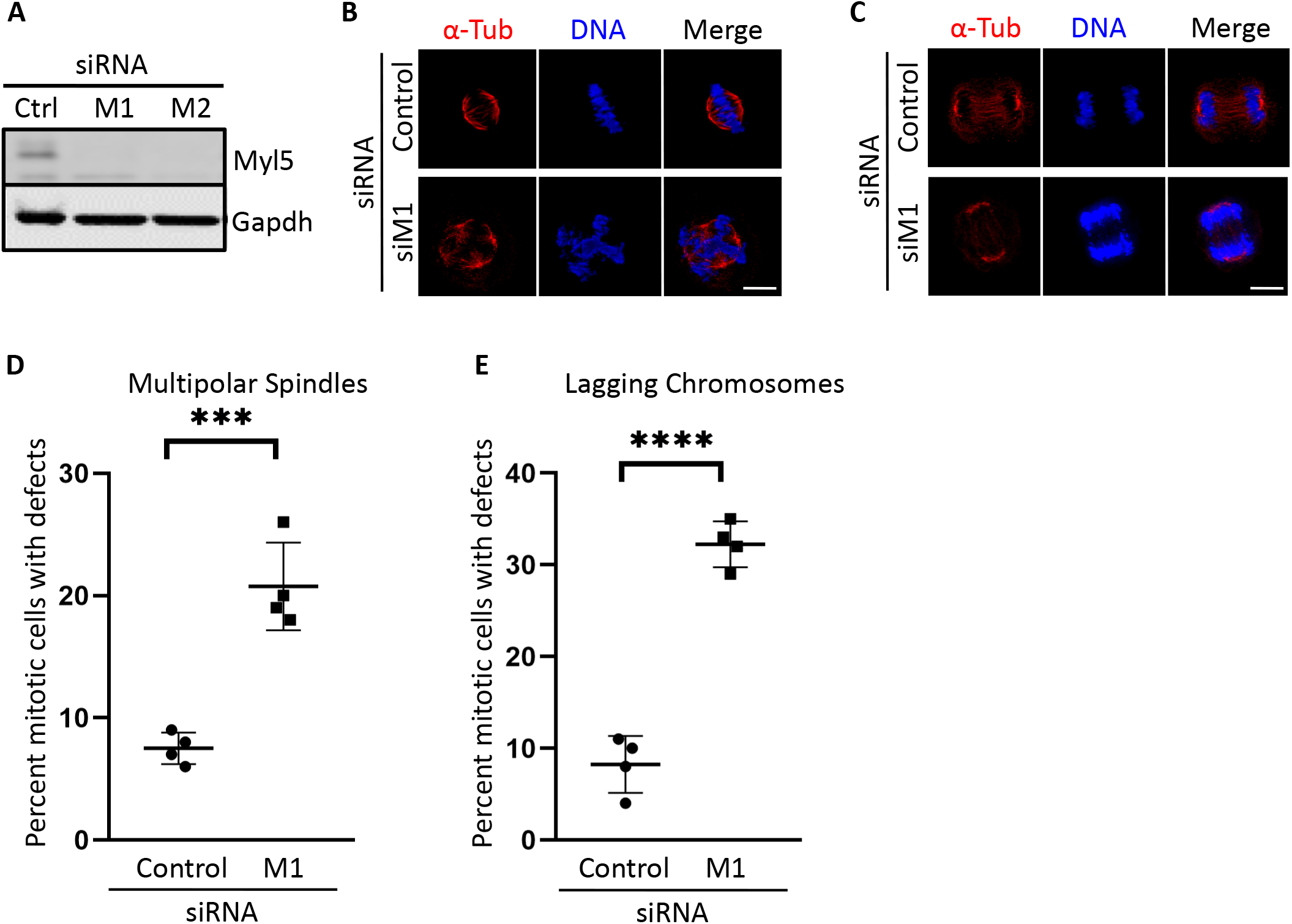
Depletion of Myl5 leads to spindle assembly and cell division defects. (**A**) siRNA knockdown of Myl5 protein levels. Immunoblot analysis showing that siRNA oligonucleotides targeting *MYL5* (M1 and M2) expression deplete Myl5 protein levels in HeLa cells compared to non-targeting control siRNA (siCtrl). (**B and C**) Immunofluorescence microscopy of HeLa cells treated with siCtrl or siM1 for 72 hours, fixed, and stained with Hoechst 33342 DNA dye and anti-a-Tubulin antibodies. Note that siM1-treated cells display multipolar spindles in prometaphase (**B**) and lagging chromosomes (**C**) in anaphase. Scale bar indicates 5μm. (**D and E**) Quantitation of the percent mitotic cells with multipolar spindles (**D**) and lagging chromosomes (**E**) in siCtrl or siM1 treated cells. Data represent the average ± SD of four independent experiments, 100 cells counted for each. *** indicates a p value =.0004 and **** a p value <.0001.

### The levels of Myl5 affect the timing of cell division

Next, we asked if the overall time to cell division was affected by the depletion of Myl5. HeLa cells were treated with non-targeting control siRNA (siCtrl) or siRNAs targeting Myl5 (siM1 and siM3) for 48 hours, synchronized in G1/S with thymidine treatment, and released in media containing the cell permeable DNA specific stain SiR-DNA (visible in the far-red channel). Five-hours post release live cells were imaged at 20X magnification at five-minute intervals for 18 hours. Movies were then analyzed and the time from nuclear DNA condensation to nuclear separation was quantified. This analysis showed that depletion of Myl5 led to a significant increase in the time that cells spent in mitosis with the average time from nuclear DNA condensation to nuclear separation for siM1= 55±27.4 (p<.0001) and siM3 = 46±22 (p=.0288) compared to siCtrl= 38.9±21.6 (**Fig. 4A and B and Movies S1-S3**). Next, we asked if Myl5 overexpression could shorten the duration of cell division. Indeed, HeLa cells that were treated with GFP-Myl5 cDNA progressed faster from nuclear DNA condensation to nuclear separation (32.18±7.5 minutes, p<.0001) compared to cells that had been treated with control GFP cDNA (55.69±35.45 minutes) (**Fig. 4C and D and Movies S4-S5**). These results indicated that depletion of Myl5 led to a slowed progression through mitosis and that overexpression of Myl5 led to a faster mitosis.

**Figure 4.**
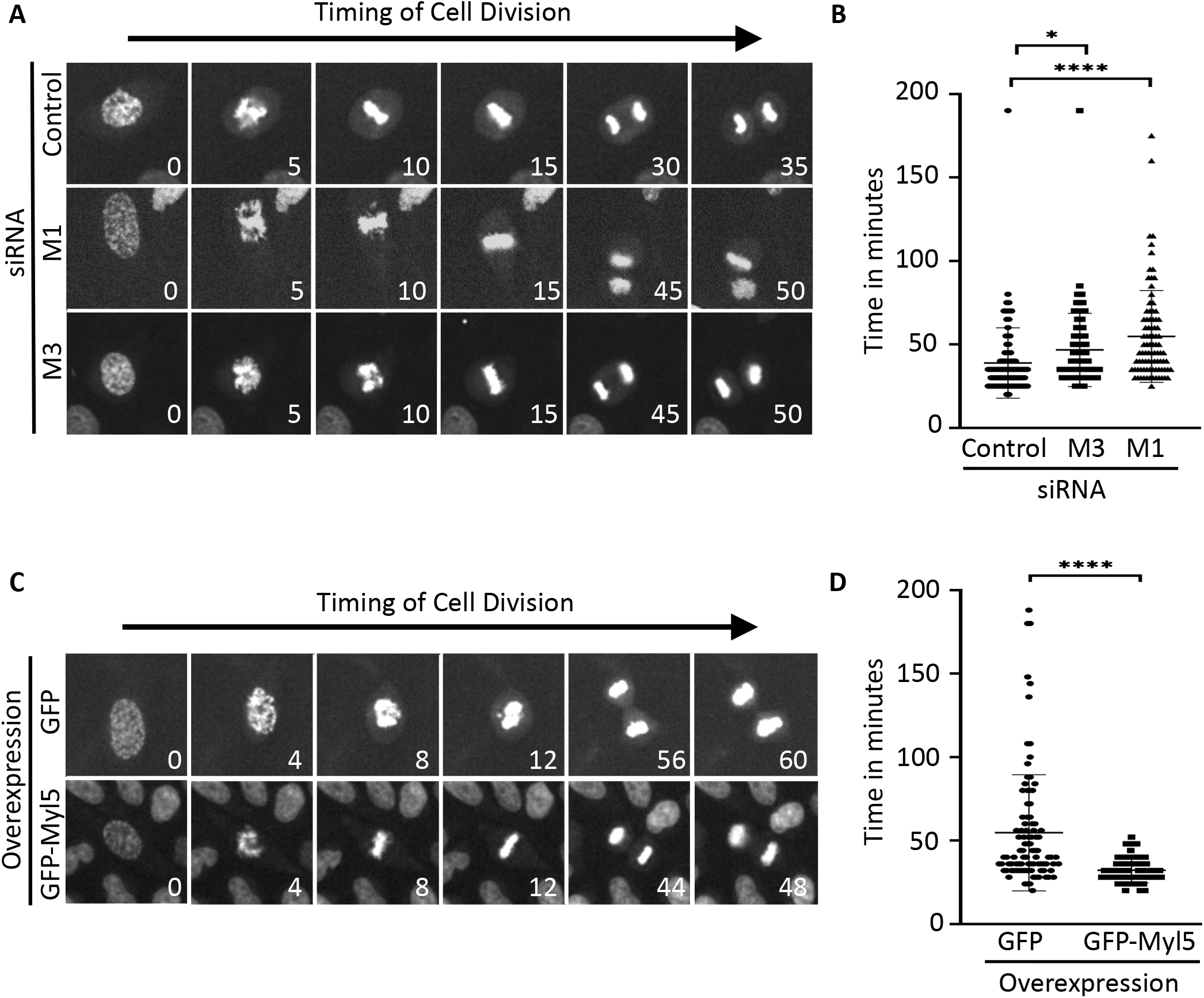
Modulation of Myl5 levels affects the time to cell division. (**A**) Live-cell time-lapse microscopy of HeLa cells treated with siCtrl or siM1 or siM3 for 72 hours. Cells were then synchronized in G1/S with thymidine treatment for 18 hours, released, and imaged by staining the cells with SiR-DNA stain at five hours post-release. The indicated times are in minutes. See supplemental Movies S1-S3. (**B**) Quantitation of the time cells spend in mitosis from DNA condensation to chromosome separation. Y-axis indicates time in minutes. X-axis indicates the siRNA treatments. Data represent the average ± SD of three independent experiments, 90 cells counted for each. * indicates p value =.0288 and **** p value <.0001. (**C**) Same as in **A**, except that cells were treated with GFP or GFP-Myl5 cDNA for 16 hours prior to live-cell time-lapse imaging. (**D**) Quantitation of the time cells spend in mitosis from DNA condensation to chromosome separation. Y-axis indicates time in minutes. X-axis indicates the cDNA treatments. Data represent the average ± SD of three independent experiments, 90 cells counted for each. **** indicates p value <.0001. See supplemental Movies S4-S5.

### Myl5 colocalizes with MYO10 to the spindle poles

Due to the ability of MYO10 to localize to the spindle poles in early mitosis and its functional importance in ensuring the fidelity of spindle assembly, chromosome congression, chromosome segregation, and cell division^14^, we asked if Myl5 and MYO10 shared a similar localization during mitosis. The LAP-Myl5 cell line was used to express GFP-Myl5 and cells were fixed, stained with Hoechst 33342 DNA dye, and anti-a-Tubulin, anti-MYO10, and anti-GFP antibodies and imaged by immunofluorescence microscopy. Indeed, the Myl5 and MYO10 localization signals overlapped at the spindle poles throughout mitosis (**Fig. 5A**). Together, these data indicated that Myl5 and MYO10 colocalize to the spindle poles during mitosis.

**Figure 5.**
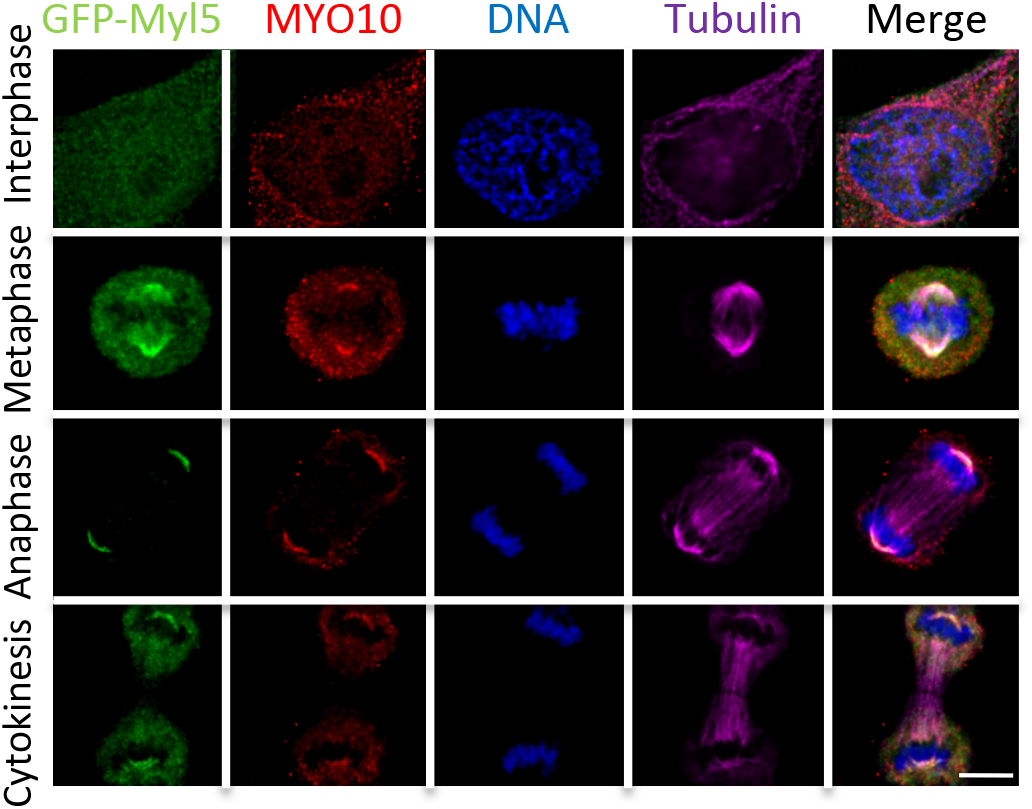
Myl5 colocalizes with MYO10 at mitotic spindle poles during mitosis. (**A and B**) The LAP (GFP-TEV-S-Peptide)-tagged-Myl5 HeLa inducible stable cell line was treated with Dox for 16 hours to express GFP-Myl5 and cells were fixed, stained with Hoechst 33342 DNA dye, and anti-a-Tubulin and anti-MYO10 antibodies and imaged by immunofluorescence microscopy. Images show the cell cycle subcellular localization of GFP-Myl5 and MYO10 during interphase, metaphase, anaphase and cytokinesis. Note that GFP-Myl5 colocalizes with MYO10 at the mitotic spindle poles during mitosis. Bar indicates 5μm.

## DISCUSSION

Although actin and the unconventional myosin MYO10 had been implicated in ensuring the fidelity of mitotic spindle assembly and cell division, the role of myosin RLCs during cell division remained unknown. Here, we have determined that Myl5 is a novel and important factor necessary for proper cell division. Myl5 dynamically localized to the spindle poles and spindle microtubules during early mitosis and depletion of Myl5 led to mitotic spindle defects, errors in chromosome congression and segregation, and a slowed progression through mitosis. These Myl5 depletion phenotypes were similar to those reported upon MYO10 depletion^14^, albeit less severe. Interestingly, the Myl5 immunofluorescence signal overlapped with MYO10 at the spindle poles throughout mitosis. Our results suggest that Myl5 may be a novel MYO10 light chain that is important for regulating MYO10 activity during cell division. To our knowledge, Myl5 is the first myosin RLC that has been implicated in mitotic spindle assembly. Of interest, a recent report showed that a calcium (Ca^2+^) signal localized to the centrosome and spindle poles throughout mitosis and was important for mammalian cell division^33^. Several proteins important for centrosome homeostasis, including the Centrin family^34^, localize to this region during cell division and contain EF-hand Ca^2+^-binding domains that are important to their function^28, 34^. Additionally, myosin light chains, including Myl5, contain EF hands and their ability to modulate myosin activity is influenced by Ca^2+^-binding^22, 23^. Therefore, it will be interesting to explore whether the localized availability of Ca^2+^ at the spindle poles is important for Myl5’s function in spindle assembly.

Our results showing that Myl5 overexpression induces a faster progression through mitosis (**Fig. 4C**) is intriguing, given that the *in silico MYL5* gene alteration frequency analysis showed that *MYL5* was amplified in various types of cancers including ovarian, bladder, adrenocortical, and pancreatic cancers (**Fig. 1C**). Therefore, it will be of interest to determine if the overabundance of Myl5 is contributing to the proliferation rates of these cancers. Additionally, other groups have observed elevated *MYL5* expression brain, cervical, and breast cancers and have linked *MYL5* overexpression to increased rates of cell migration, metastasis, and poor survival^25–27^. More broadly, of the ~40 myosins encoded in the human genome, at least ten (including MYO10) have been implicated in tumorigenesis^35^. Therefore, it is very likely that Myl5 has important roles outside of mitosis and that it may be regulating myosin activity to influence the cytoskeletal network during cell migration and metastasis. Molecular studies aimed at analyzing the role of Myl5 in these processes, which are key to tumorigenesis, are likely to further advance our understanding of Myl5 in cancer. We note that during interphase, myosin light chains have been implicated in regulating gene expression by binding to specific sequences within the promoter region of target genes^36, 37^ and that Myl5 has been shown to bind the promoter region of HIF-1alpha, an important factor in tumorigenesis, and regulate its expression ^25, 38^. Consistent with this function Myl5 localized to both the cytoplasm and nucleus in interphase cells (**Fig. 2A**). Therefore in addition to its cytoskeleton-related function, Myl5 may have cytoskeleton unrelated functions that contribute to tumorigenesis.

## MATERIALS and METHODS

### Cell culture

HeLa cells were grown in F12:DMEM 50:50 (Hyclone) with 10% FBS, 2mM L-glutamine and antibiotics in 5% CO_2_ at 37°C. Cells were synchronized in G1/S by treatment with 2 mM thymidine (Sigma-Aldrich) for 18-hours. The following siRNAs were used for siRNA treatments: ThermoFisher Silencer Select 4390843 (control non-targeting siRNA) and S9187 and S9188 (M1 and M2 siRNAs targeting *MYL5*); Dharmacon ON-TARGETplus D-001810-10 (control non-targeting siRNA) and J-011739-03 and J-011739-04 (M3 and M4 siRNAs targeting *MYL5*) were used as described previously.^39^ Please see Table S1 for a list of key reagents and resources used in this study and their pertinent information.

### Generation of the LAP-Myl5 inducible stable cell line

The HeLa LAP(GFP-TEV-S-Peptide)-Myl5 stable cell line was generated according to Torres et al.^31, 32^ Briefly, full-length *MYL5* (coding for amino acid residues 1-173) was cloned into pDONR221 and transferred to pGLAP1 through a Gateway reaction to generate the pGLAP1-*MYL5* vector that was transfected into HeLa Flp-In T-Rex cells to generate the LAP-Myl5 inducible stable cell line as described previously.^31, 32^

### Immunoblotting

For Myl5 cell cycle protein expression analysis, HeLa cells were synchronized in G1/S with 2 mM thymidine for 18-hours. Cells were then washed with PBS three times and twice with F12:DMEM media with 10% FBS and released into the cell cycle. Cells were harvested at the indicated time points, lysed, and protein extracts were resolved on a 4-20% SDS-PAGE and transferred to a PVDF membrane. The membranes were immunoblotted with the indicated antibodies and imaged with a LiCOR system. The same approach was used to detect Myl5 protein depletion upon siRNA treatments without the cell synchronization step. Cell extract preparation and immunoblot analyses with the indicated antibodies were as described previously.^40^

### Fixed-cell immunofluorescence microscopy and live-cell time-lapse microscopy

Fixed-cell immunofluorescence microscopy was performed as described in Gholkar et al.^40^ Briefly, non-transfected cells or cells that had been transfected with the indicated siRNAs for 48 hours were arrested in G1/S with 2 mM thymidine for 18 hours, washed, and released into fresh media for ten hours. Cells were then fixed with 4% paraformaldehyde, permeabilized with 0.2% Triton X-100/PBS, and co-stained with 0.5 μg/ml Hoechst 33342 (ThermoFisher) to visualize the DNA and the indicated antibodies. A Leica DMI6000 microscope (Leica DFC360 FX Camera, 63x/1.40-0.60 NA oil objective, Leica AF6000 software) was then used to capture the images, which were deconvolved with the Leica Application Suite 3D Deconvolution software and exported as TIFF files. For quantifying mitotic defects, the data from four independent experiments, with 100 cells counted for each, was used to quantify the average ± standard deviation (SD). For time-lapse microscopy, HeLa cells were transfected with the indicated siRNAs or cDNAs for 24 hours, arrested in G1/S with 2 mM thymidine for 18 hours, washed, and released into fresh media containing 100 nM SiR-DNA stain (Cytoskeleton Inc.). Cells were imaged live five-hours post release for 18 hours using an ImageXpress XL imaging system (Molecular Devices) with a 20x air objective at 37 °C in 5% CO_2_. Captured images were exported as a video at two frames per second using Image J and the videos were saved as AVI movies. Each frame represents a five-minute interval. For quantifying the timing of cell division, the data from three independent experiments, with 90 cells counted for each, was used to quantify the average time in minutes from DNA condensation for nuclear separation ± standard deviation (SD).

### In silico analysis of Myl5

Myl5 ortholog analysis was performed by querying OrthoDB^29^ (https://www.orthodb.org/) for Myl5 (ID: 177713at9347) at the Eutheria level. *MYL5* genomic alteration frequencies in tumor samples were analyzed using The Cancer Genome Atlas (TCGA) Pan-Cancer Atlas^30^ (https://www.cell.com/pb-assets/consortium/pancanceratlas/pancani3/index.html), which includes over 11,000 tumors from 33 most prevalent cancer forms. Query was run considering at least ten total cancer cases and results were displayed as percent alteration frequencies by cancer type.

### Antibodies

Immunofluorescence, immunoblotting, and immunoprecipitations were carried out using antibodies against: Myl5 and MYO10 (Proteintech: 14249-1-AP and 24565-1-AP); Pericentrin (Novus Biologicals: NB-100-68277); GFP (Abcam: ab290); Gapdh, S-tag (GeneTex: GTX128060 and GTX100118); a-Tubulin (Serotec: MCAP77); Cyclin B (Santa Cruz: sc-245). Centrin antibodies were a gift from J. Salisbury and NUMA and TPX2 antibodies were gifts from D. Compton. Secondary antibodies conjugated to FITC, Cy3, and Cy5 were from Jackson Immuno Research and those conjugated to IRDye 680 and IRDye 800 were from LI-COR Biosciences.

## ACKNOWLEDGMENTS

This material is based upon work supported by the National Science Foundation under Grant Number MCB1243645 to J.Z.T., any opinions, findings, and conclusions or recommendations expressed in this material are those of the authors and do not necessarily reflect the views of the National Science Foundation. This work was supported in part by a grant to The University of California, Los Angeles from the Howard Hughes Medical Institute through the James H. Gilliam Fellowships for Advanced Study program (E.F.V). This work was also supported by an NSF Louis Stokes Alliances for Minority Participation Bridge to the Doctorate Fellowship and Cota Robles Fellowship to I.R.

