## Supplementary Material for "The myosin regulatory light chain Myl5 localizes to mitotic spindle poles and is required for proper cell division"

### SUPPLEMENTAL FIGURES

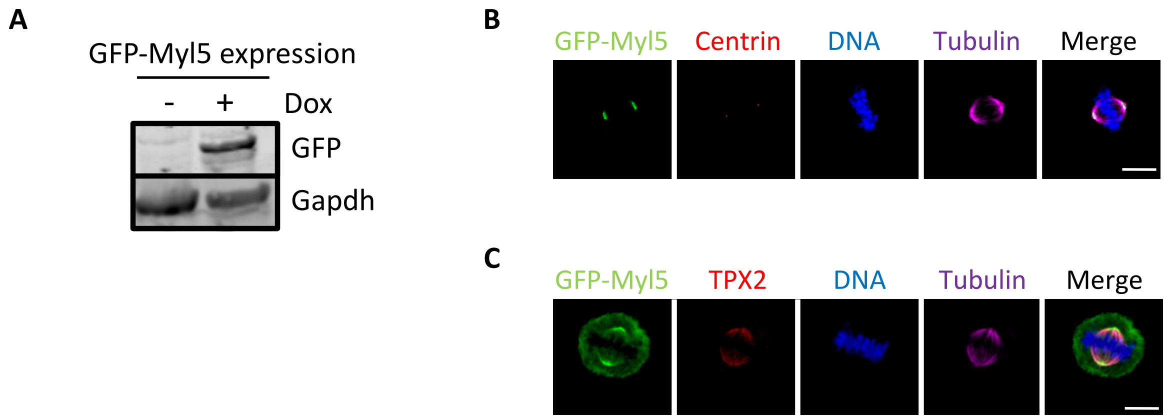


**Supplemental Figure S1.** Expression of GFP-Myl5 and its localization to mitotic spindle poles during mitosis. (**A**) Immunoblot analysis of protein extracts from the Dox inducible LAP (GFP-TEV-S-Peptide)-tagged-Myl5 stable cell line in the absence (-) or presence (+) of Dox. Blots were probed with anti-GFP or anti-Gapdh antibodies. Immunoblot shows that GFP-Myl5 is expressed only when cells are treated with Dox. (**B** and **C**) The LAP (GFP-TEV-S-Peptide)-tagged-Myl5 HeLa inducible stable cell line was treated with Dox for 16 hours to express GFP-Myl5 and cells were fixed, stained with Hoechst 33342 DNA dye and anti-α-Tubulin, anti-GFP, anti-Centrin (**B**), and anti-TPX2 (**C**) antibodies and imaged by immunofluorescence microscopy. Images show the localization of GFP-Myl5 during metaphase in relation to Centrin and TPX2. Bars indicate 5μm.

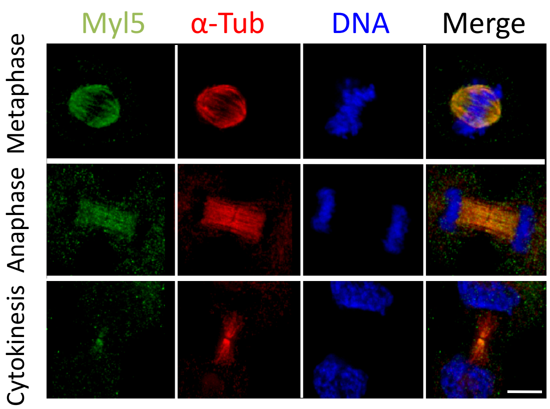


**Supplemental Figure S2.** Endogenous Myl5 localizes to the spindle and spindle poles during mitosis. HeLa cells were fixed, stained with Hoechst 33342 DNA dye and anti-α-Tubulin and anti-Myl5 antibodies and imaged by immunofluorescence microscopy. Images show the localization of Myl5 during metaphase, anaphase, and cytokinesis. Bar indicates 5μm.

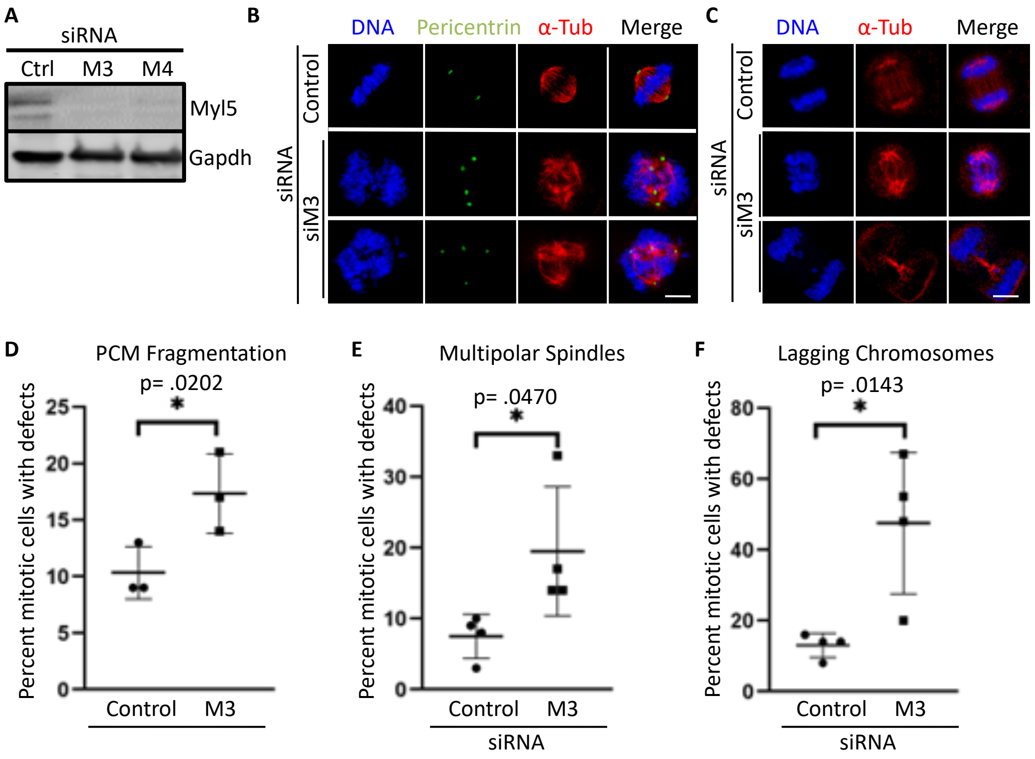


**Supplemental Figure S3.** Depletion of Myl5 leads to spindle assembly and cell division defects. (**A**) siRNA knockdown of Myl5 protein levels. Immunoblot analysis showing that siRNA oligonucleotides targeting *MYL5* (M3 and M4) expression deplete Myl5 protein levels in HeLa cells compared to non-targeting control siRNA (siCtrl). (**B** and **C**) Immunofluorescence microscopy of HeLa cells treated with siCtrl or siM3 for 48 hours, fixed, and stained with Hoechst 33342 DNA dye and anti-α-Tubulin and anti-Pericentrin (**B**) antibodies. Note that siM3-treated cells display multipolar spindles in prometaphase (**B**), Pericentrin (marker for pericentriolar material- PCM) fragmentation with >2 foci (**B**), and lagging chromosomes (**C**) in anaphase. Scale bar indicates 5µm. (**D-F**) Quantitation of the percent mitotic cells with PCM fragmentation (**D**), multipolar spindles (**E**) and lagging chromosomes (**F**) in siCtrl or siM3 treated cells. Data represent the average ± SD of 4 independent experiments, 100 cells counted for each. * indicates a p value <.05 as indicated above each asterisk.

### SUPPLEMENTAL TABLES

#### Supplemental Table S1. Reagents and Resources

| REAGENT or RESOURCE | SOURCE | IDENTIFIER |
|---|---|---|
| Antibodies | | |
| Chicken polyclonal anti-GFP | Abcam | Cat# ab13970; RRID: AB_300798 |
| Rabbit polyclonal anti-HA | Proteintech | Cat# 51064-2-AP; RRID: AB_11042321 |
| Rat monoclonal anti-α-Tubulin (clone YOL1/34) | Bio-Rad | Cat# MCA78G; RRID: AB_325005 |
| Rabbit polyclonal anti-MYO10 | Proteintech | Cat# 24565-1-AP |
| Rabbit polyclonal anti-Pericentrin | Novus | Cat# NB100-68277; RRID:AB_1109761 |
| Rabbit polyclonal anti-GAPDH | Genetex | Cat# GTX100118; RRID:AB_1080976 |
| Rabbit polyclonal anti-MYL5 | Proteintech | Cat# 14249-1-AP; RRID:AB_2266762 |
| Mouse monoclonal anti-Cyclin B1 | Santa Cruz | Cat# sc-245; RRID:AB_627338 |
| Rabbit polyclonal anti-NUMA | Gift from D. Compton | N/A |
| Rabbit polyclonal anti-TPX2 | Gift from D. Compton | N/A |
| Rabbit polyclonal anti-Centrin | Gift from J. Salisbury | N/A |
| Donkey polyclonal anti-Human IgG (H+L), Fluorescein (FITC) AffiniPure | Jackson ImmunoResearch Labs | Cat# 709-095-149; RRID: AB_2340514 |
| Donkey polyclonal anti-Rat IgG (H+L), Cy3 AffiniPure | Jackson ImmunoResearch Labs | Cat# 712-165-153; RRID: AB_2340667 |
| Donkey polyclonal anti-Human IgG (H+L), Cy5 AffiniPure | Jackson ImmunoResearch Labs | Cat# 709-175-149; RRID: AB_2340539 |
| Donkey polyclonal anti-Chicken IgY (IgG) (H+L), Fluorescein (FITC) AffiniPure | Jackson ImmunoResearch Labs | Cat# 703-095-155; RRID: AB_2340356 |
| Donkey polyclonal anti-Rabbit IgG (H+L), Cy3 AffiniPure | Jackson ImmunoResearch Labs | Cat# 711-165-152; RRID: AB_2307443 |
| Donkey polyclonal anti-Rabbit IgG (H+L), Fluorescein (FITC) AffiniPure | Jackson ImmunoResearch Labs | Cat# 711-095-152; RRID: AB_2315776 |
| Donkey polyclonal anti-Mouse IgG (H+L), Fluorescein (FITC) AffiniPure | Jackson ImmunoResearch Labs | Cat# 715-095-151; RRID: AB_2335588 |
| Donkey polyclonal anti-Mouse IgG (H+L), Cy3 AffiniPure | Jackson ImmunoResearch Labs | Cat# 715-165-151; RRID: AB_2315777 |
| Donkey polyclonal anti-Goat IgG (H+L), IRDye 680RD | LI-COR Biosciences | Cat# 926-68074; RRID: AB_10956736 |
| Donkey polyclonal anti-Mouse IgG (H+L), IRDye 680RD | LI-COR Biosciences | Cat# 926-68072; RRID: AB_10953628 |
| Donkey polyclonal anti-Mouse IgG (H+L), IRDye 800CW | LI-COR Biosciences | Cat# 926-32212; RRID: AB_621847 |
| Donkey polyclonal anti-Chicken IgG (H+L), IRDye 800CW | LI-COR Biosciences | Cat# 926-32218; RRID: AB_1850023 |
| Donkey polyclonal anti-Rabbit IgG (H+L), IRDye 680RD | LI-COR Biosciences | Cat# 926-68073; RRID: AB_10954442 |
| Donkey polyclonal anti-Rabbit IgG (H+L), IRDye 800CW | LI-COR Biosciences | Cat# 926-32213; RRID: AB_621848 |
| Chemicals, Peptides, and Recombinant Proteins | | |
| Paclitaxel | Sigma-Aldrich | Cat# T7191; CAS:33069-62-4 |
| Nocodazole | Sigma-Aldrich | Cat# 1404; CAS:31430-18-9 |

| | | |
|---|---|---|
| Thymidine | Sigma-Aldrich | Cat# T1895; CAS:50-89-5 |
| Doxycycline | Sigma-Aldrich | Cat# D9891; CAS:24390-14-5 |
| Halt Protease Inhibitor Cocktail | Thermo Fisher Scientific | Cat# 87786 |
| Hoechst 33342 | Thermo Fisher Scientific | Cat# H1399; CAS:23491-52-3 |
| ProLong Gold Antifade Mountant | Thermo Fisher Scientific | Cat# P36934 |
| Lipofectamine RNAiMAX | Thermo Fisher Scientific | Cat# 13778150 |
| FuGENE HD | Promega | Cat# E2311 |
| FuGene 6 | Promega | Cat# E2691 |
| S-protein Agarose | Millipore Sigma | Cat# 69704-4 |
| Anti-FLAG M2 magnetic beads | Sigma-Aldrich | Cat# M8823 |
| Dynabeads MyOne Streptavidin C1 | Thermo Fisher Scientific | Cat# 65002 |
| Biotin | Sigma | Cat# B4501 |
| Affi-Prep Protein A beads | Bio-Rad | Cat# 156-0005 |
| GFP-Trap Dynabeads | Chromotek | Cat# gtd-20 |
| Critical Commercial Assays | | |
| SP6 TnT Quick Coupled Transcription/Translation System | Promega | Cat# L2080 |
| PureYield Plasmid Miniprep System | Promega | Cat# A1222 |
| QIAprep Spin Miniprep Kit | QIAGEN | Cat# 27106 |
| PureYield Plasmid Midiprep System | Promega | Cat# A2495 |
| Experimental Models: Cell Lines | | |
| Human: HeLa cells | ATCC | Cat# CCL-2; RRID: CVCL_0030 |
| Human: HCT116 cells constitutively expressing GFP-H2B | Gift from P. Jackson | N/A |
| Human: HeLa Flp-In T-Rex cells | Gift from S. Taylor | N/A |
| Oligonucleotides | | |
| Silencer™ Select siRNA targeting *MYL5* | Thermo Fisher Scientific | Cat# 4392420; siRNA ID: s9187 |
| Silencer™ Select siRNA targeting *MYL5* | Thermo Fisher Scientific | Cat# 4392420; siRNA ID: s9188 |
| Silencer™ Select Negative Control siRNA | Thermo Fisher Scientific | Cat# 4390843 |
| ON-TARGETplus Human *MYL5* siRNA | Dharmacon | Cat# J-011739-03 |
| ON-TARGETplus Human *MYL5* siRNA | Dharmacon | Cat# J-011739-04 |
| ON-TARGETplus Non-targeting Control Pool | Dharmacon | Cat# D-001810-10 |
| Recombinant DNA | | |
| *MYL5* cDNA | GenScript | Clone ID: OHu30367 |
| pDONR221-Myl5 | This paper | N/A |
| pGLAP1-Myl5 | This paper | N/A |
| pCS2-HA-Myl5 | This paper | N/A |
| pCS2-FLAG-Myl5 | This paper | N/A |
| pGBioID2-Myl5 | This paper | N/A |

| | | |
|---|---|---|
| pDONR221-MYO10 | This paper | N/A |
| pGLAP1-MYO10 | This paper | N/A |
| EGFPC1-hMYO10 | Addgene | Clone ID: 47608 |
| pCS2-HA-MYO10 | This paper | N/A |
| pCS2-FLAG-MYO10 | This paper | N/A |
| pENTR223-TPX2 | DNASU | Clone ID: HsCD00505542 |
| pDONR221-TPX2 | This paper | N/A |
| pGLAP1-TPX2 | This paper | N/A |
| pCS2-HA-TPX2 | This paper | N/A |
| pCS2-FLAG-TPX2 | This paper | N/A |
| pDONR221-GFP | This paper | N/A |
| pGLAP1-GFP | This paper | N/A |
| pCS2-HA-GFP | This paper | N/A |
| pCS2-FLAG-GFP | This paper | N/A |
| pDONR221 | Thermo Fisher Scientific | Cat# 12536017 |
| Software and Algorithms | | |
| GraphPad Prism 8 | GraphPad | RRID: SCR_002798 |
| ImageJ | NIH ImageJ | https://imagej.nih.gov/ij/index.html RRID:SCR_003070 |

### SUPPLEMENTAL MOVIES

**Supplemental Movie S1.** Live cell time-lapse microscopy movie of a siControl-treated HeLa cell undergoing cell division. Cells were arrested with 2 mM thymidine for 18 hours, washed and released into fresh media. Cells were imaged at 5 hours post release for 18 hours every five minutes using an ImageXpress XL microscope at 20X magnification, 37 $^{0}$C, and 5% $CO_2$. Images were converted to an AVI movie format. Each frame represents a five-minute interval.

**Supplemental Movie S2.** Live cell time-lapse microscopy movie of a siM1-treated HeLa cell undergoing cell division, as described for supplemental Movie S1.

**Supplemental Movie S3.** Live cell time-lapse microscopy movie of a siM3 treated HeLa cell undergoing cell division, as described for supplemental Movie S1.

**Supplemental Movie S4.** Live cell time-lapse microscopy movie of a HeLa cell transfected with GFP cDNA undergoing cell division. Cells were arrested with 2 mM thymidine for 18 hours, washed and released into fresh media. Cells were imaged at 5 hours post release for 18 hours every five minutes using an ImageXpress XL microscope at 20X magnification, 37 $^{0}$C, and 5% $CO_2$. Images were converted to an AVI movie format. Each frame represents a five-minute interval.

**Supplemental Movie S5.** Live cell time-lapse microscopy movie of a HeLa cell transfected with GFP-Myl5 cDNA undergoing cell division, as described for supplemental Movie S4.
